# What is the role of the environment in the emergence of novel antibiotic resistance genes? – A modelling approach

**DOI:** 10.1101/2021.04.04.438392

**Authors:** Johan Bengtsson-Palme, Viktor Jonsson, Stefanie Heß

**Affiliations:** Department of Infectious Diseases, Institute of Biomedicine, The Sahlgrenska Academy, University of Gothenburg, Guldhedsgatan 10, SE-413 46, Gothenburg, Sweden; Centre for Antibiotic Resistance research (CARe) at University of Gothenburg, Gothenburg, Sweden; Integrated Science Lab, Department of Physics, Umeå University, SE-901 87 Umeå, Sweden; Institute of Microbiology, Technische Universität Dresden, Zellescher Weg 20b, 01847 Dresden, Germany

## Abstract

It is generally accepted that intervention strategies to curb antibiotic resistance cannot solely focus on human and veterinary medicine but must also consider environmental settings. While the environment clearly has a role in the transmission of resistant bacteria, it is less clear what role it plays in the emergence of novel types of resistance. It has been suggested that the environment constitutes an enormous recruitment ground for resistance genes to pathogens, but the extent to which this actually happens is unknown. In this study, we built a model framework for resistance emergence and used the available quantitative data on the relevant processes to identify the steps which are limiting the appearance of antibiotic resistance determinants in human or animal pathogens. We also assessed the effect of uncertainty in the available data on the model results. We found that in a majority of scenarios, the environment would only play a minor role in the emergence of novel resistance genes. However, the uncertainty around this role is enormous, highlighting an urgent need of more quantitative data to understand the role of the environment in antibiotic resistance development. Specifically, more data is most needed on the fitness costs of antibiotic resistance gene (ARG) carriage, the degree of dispersal of resistant bacteria from the environment to humans, but also the rates of mobilization and horizontal transfer of ARGs. Quantitative data on these processes is instrumental to determine which processes that should be targeted for interventions to curb development and transmission of resistance.

## Introduction

Antibiotic resistance is a globally growing health threat, which is projected to take more lives than all forms of cancer combined by 2050 if it cannot be controlled ^1^. Nowadays, there is a general consensus that intervention strategies should consider not only human and veterinary medicine but also take into account the environment, in what is referred to as a one health approach ^2^. While surveillance programs and careful reduction of the use of antibiotics in clinical and agricultural settings started years ago, we are still only starting to understand the role of the environment in the development and dissemination of antibiotic resistance ^3–5^. Based on monitoring data on the abundance and diversity of antibiotic resistance genes (ARGs), we know conceptually which the dispersal routes between and within the three compartments (humans, farmed animals and the external environment) are ^6–8^. We also have a reasonably clear picture of what processes result in recruitment of resistance factors from environmental bacteria to human pathogens ^9^. However, much remains to be done in terms of quantifying the relative importance of these processes, which is indispensable knowledge for effective strategies to minimize the spread and development of antibiotic resistance ^10^.

It has been proposed that the environment could have three main roles in resistance development ^9^. Firstly, it enables the transfer of resistance genes between environmental, human and animal associated bacteria. Secondly, it constitutes a reservoir or intermediate habitat for resistant bacteria and resistance genes. Finally, it can play a crucial role in the evolution of novel resistance factors, as it provides an arena for selection of resistance combined with an enormous source of genetic diversity from which bacteria can recruit resistance genes ^8,9^. Newly emerged resistance genes from the environment can subsequently be disseminated into the human and farmed animals compartments, where they have the potential to cause severe health threats. While this conceptual role of the environment is well described, its actual importance in resistance development is so far unknown, largely due to a lack of quantitative data ^10,11^. Particularly, the probability of emergence of novel resistance determinants in the environment and their transfer rates to human pathogens are completely unknown this far ^11^. This means that it becomes very hard to rank risk scenarios and prioritize between possible interventions targeting the emergence of novel resistance factors. The primary aim of this study is to identify the steps within the chain of processes which are limiting the appearance of antibiotic resistance determinants in human pathogens. To address this, we set up a model framework and quantified the dependencies of variables given estimates for upper and lower bounds for the process rates available in literature. Furthermore, we assess how the uncertainty in the available data affects certain parameters in the model, thereby highlighting where the most urgent needs for quantitative data exist.

## Methods

### Implementation of the model

We consider two different scenarios in our model. In the first scenario, ARGs that later end up in human pathogens already pre-existed in bacteria in some setting at the start of the antibiotic era (around 70 years ago), and only needed to be, in some way, transferred to human pathogens (Table 1). In the second model scenario, we assume that ARGs did not have a resistance function at the start of the modelled time period but needed to first emerge as resistance factors. We will refer to the latter scenario as the “emergence model” and the first scenario as the “pre-existing model”.

**Table 1.** Considered pathways and the corresponding equations in the two models

| Description of pathway | Equation (main model) | Equation (emergence model) |
| --- | --- | --- |
| Appearance directly on a mobile genetic element in a human pathogen | $E1(t) = 10^{30} * Pph * Pm * E * S^t$ | $E1(t) = 10^{30} * Pph * Pm * E * t * S^{t/2}$ |
| Appearance in non-pathogenic human bacteria, mobilized and transferred to human pathogens | $E2(t) = 10^{30} * Ph * E * MH * t * S^t$ | $E2(t) = 10^{30} * Ph * E * t * MH * t/2 * S^{t/2}$ |
| Appearance on a mobile genetic element in non-pathogenic human-associated bacteria | $E3(t) = 10^{30} * Ph * Pm * E * H * t * S^t$ | $E3(t) = 10^{30} * Ph * Pm * E * t * H * t/2 * S^{t/2}$ |
| Appearance in pathogens in environment and disseminated to humans | $E4(t) = 10^{30} * Pp * Pm * E * S^t * D * t$ | $E4(t) = 10^{30} * Pp * Pm * E * t * S^{t/2} * D * t/2$ |
| Appearance in environmental bacteria, mobilized, transferred to pathogens and disseminated to humans | $E5(t) = 10^{30} * E * MH * t * S^t * D * t/2$ | $E5(t) = 10^{30} * E * t * MH * t/2 * S^{t/2} * D * t/4$ |
| Appearance on a mobile genetic element in environmental bacteria, transferred to pathogens and disseminated to humans | $E6(t) = 10^{30} * Pm * E * H * t * S^t * D * t/2$ | $E6(t) = 10^{30} * Pm * E * t * H * t/2 * S^{t/2} * D * t/4$ |

For process and parameter abbreviations, see Table 2. $t$ is for time in days. $10^{30}$ represents the approximate total number of bacteria on earth.

The endpoint of the proposed models is a mobilized ARG observed in a human pathogen. Several different chains of events may lead from the first appearance of an ARG to its occurrence in a pathogenic strain (Fig.1). These combinations of events are summarized in the model as six different pathways. First, an ARG may have existed directly on a mobile genetic element in a human pathogen living in a human being (E1). Secondly, an ARG could have already existed on the chromosome of a human commensal bacteria. After mobilization, the ARG could then be horizontally transferred to a human pathogen (E2). Thirdly, an ARG could originate from a mobile genetic element in a commensal species belonging to the human microbiome. In this scenario, it would then be directly transferrable into a human pathogen via HGT (E3). These first three processes all target pre-existence of ARGs in bacteria directly associated with humans. However, it is also plausible that ARGs may have originated in bacteria living in the environment and subsequently been disseminated to humans. The simplest case would be an ARG that originated from a human pathogen present in the environment. It could then be disseminated between the compartments before ending up in humans (E4). Alternatively, a chromosomally encoded ARG could have existed among environmental bacteria that are not pathogenic. Subsequently, that ARG needs to be mobilized onto a plasmid or as a transposon, spread via horizontal gene transfer (HGT, referred to with the letter ‘H’ in the model) and be transferred to human pathogens. Humans might pick up the bacterium harbouring the ARG, for instance, with food or while swimming (E5). An ARG could also have pre-existed on a mobile genetic element in an environmental bacterium. Via transformation, transduction or conjugation (H), it could then get transferred to a human pathogen, which can subsequently be taken up by humans (E6).

**Fig.1.**
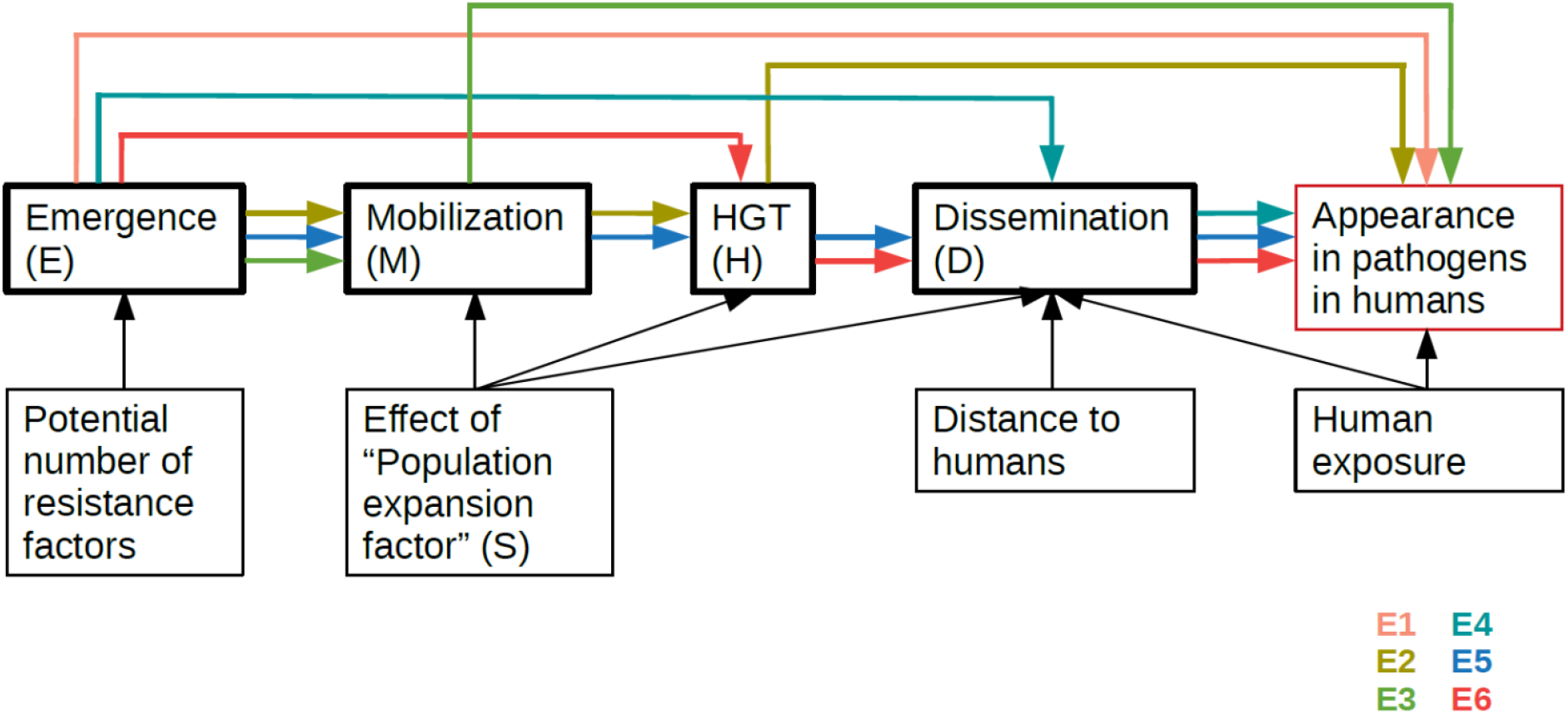
Overview of the model framework. Processes are framed in bold. Arrows display the influence of different parameters on the processes. Coloured arrows represent the pathways included in the model (E1 to E6).

Time plays an important role in all processes relating to the spread and maintenance of ARGs. For mobilization (M), horizontal gene transfer (H) and the dissemination processes (D), it is important that the microbe carrying the gene survives. Many human pathogenic species are known to have a rather short lifetime outside of their host ^12,13^. Furthermore, the fitness effect of ARG carriage on population size over time is cumulative, affecting the population expansion rate (S) exponentially, in contrast to the other model parameters. The dissemination efficiency is additionally influenced by the distance to humans in terms of space and time. Human exposure is a critical factor for the dissemination as well as the appearance of the ARG in pathogens in the human microbiome. Humans are exposed to bacteria in many different ways through which uptake via food seems to be the most important one in terms of bacterial exposure per day ^14^.

Each of the six pathways shown in Fig. 1 were expressed as equations (Table 1) with parameters defined in Table 2. The interpretation of each equation is the number of mobilized ARGs in pathogens at time t contributed by the corresponding pathway. Both models (the main “pre-existing” model and the “emergence” model) were implemented in R as follows, only differing in the meaning and range of the parameter E (Table 2). Each parameter was randomly selected from its log10 transformed range (see Table 2) under a uniform distribution. For each set of randomly selected parameters, the result of the equations for the different considered processes were calculated (Table 1). Next, the total appearance was calculated based on these sub-equations according to this formula:

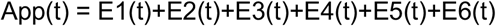

where E1 to E6 each represent the contribution of ARGs by a particular process, expressed as total contributed ARGs over the entire model timeframe, and t is the time in days, resulting in the total appearance (App) representing the total observed ARGs at time t. The true total appearance was approximated from the ResFinder database ^15^, which depending on how gene variants are counted held on the order of 700 to 2200 unique ARGs in September 2019. By assuming that these genes would all have appeared in the past 70 years since antibiotics were widely introduced, that gives us an expected appearance rate of around 9 to 29 ARGs per year, or 0.025 to 0.079 per day (Table 2). If the modelled total appearance fell within the expected interval, the model parameters were saved. This procedure was iterated until 10,000 valid sets of model parameters had been obtained for each modelled time point and set of parameter ranges.

**Table 2.** Estimates for the respective process rates and parameters used in the model

| Abbreviation | Meaning | Unit | Literature values |  |  | Model boundaries |  |
| --- | --- | --- | --- | --- | --- | --- | --- |
|  |  |  | Lower bound | Upper bound | Median | Lower bound | Upper bound |
| E | Proportion of bacteria in which a given ARG exists at the model start<br>or<br>Emergence rate of a novel ARG (in emergence model) | -<br>Events/cell/day | Unknown<br>Unknown | | | $10^{-30}$<br>$10^{-40}$ | 1<br>1 |
| App | First detected appearance of a novel ARG in a human pathogen in a human | Events/day | 0.025 | 0.079 | N/A | 0.025 | 0.079 |
| M | Mobilization of a chromosomally encoded ARG onto a plasmid, as a transposon or other conjugative element | Events/cell/day | Difficult to disentangle from horizontal transfer based on current experimental evidence, see MH |  |  |  |  |
| H | Horizontal gene transfer (conjugation) | Events/cell/day | $2.4 * 10^{-10}$ | $5.8 * 10^{-2}$ | $3.0 * 10^{-3}$ | $10^{-11}$ | $10^{-1}$ |
| D | Dissemination from the environment to humans | Events/cell/day | $10^{-14}$ | $10^{-11}$ | N/A | $10^{-15}$ | $10^{-10}$ |
| Pph | Fraction of all bacterial cells which are human pathogens and living in/on humans | - | $\sim 10^{-10}$ ($\sim 10^{13}$ bacterial cells per human * $\sim 10^{10}$ humans worldwide * $\sim 10^{-3}$ bacteria living in humans pathogenic / $\sim 10^{30}$ bacterial cells on the world) | | | $10^{-12}$ | $10^{-8}$ |
| Ph | Fraction of all bacterial cells that live in/on humans of total bacterial cells on the world | - | $\sim 10^{-7}$ ($\sim 10^{13}$ bacterial cells per human * $\sim 10^{10}$ humans worldwide / $\sim 10^{30}$ bacterial cells on the world) | | | $10^{-8}$ | $10^{-6}$ |
| MH | Mobilization of a chromosomally encoded ARG and transfer into another strain | Events/cell/day | $4 * 10^{-10}$ | $1.5 * 10^{-4}$ | $\sim 5.67 * 10^{-9}$ | $10^{-15}$ | $10^{-2}$ <sup>a</sup> |
| S | Population expansion rate (1 corresponds to no change in population size) | day<sup>-1</sup> | Unknown |  |  | 0.9 | 1.1 |
| Pm | Probability that a novel ARG emerges on a plasmid | - | 0.002 (~ 7% of all bacterial cells carry a conjugative plasmid, most plasmids have ~ 50-100 genes; about 3% of the total bacterial genome) |  |  | 0.0001 | 0.01 |
| Pp | Fraction of human pathogenic cells (in all compartments) of bacterial cells on the world | - | $10^{-9}$, must be larger than Pph, but how much larger is unknown | | | $10^{-11}$ | $10^{-7}$ |

<sup>a</sup> MH is also restricted to be < H.

## Results

### Defining the probability boundaries for antibiotic resistance development

In order to populate the model, measured values for the individual parameters were taken from literature (Supplementary Tables 1 and 2), and the upper and lower boundaries were fed into the model (Table 2). For the rates of horizontal gene transfer (H) most studies were found for the conjugative transfer of genes. Data for both intra- and interspecies transfer of various naturally occurring and artificially constructed plasmids are available. These have been included as a first approximation in the model, bearing in mind that this is laboratory data and that their transferability to the respective ecosystems has been little studied so far. At this time, we have limited knowledge concerning the role of transduction and transformation for the distribution of resistance genes in the respective ecosystems, although recent studies have suggested that they may be important in the transfer of ARGs between bacteria ^16^. Due to the lack of quantitative data, these two processes were not considered for the estimation of the parameter H.

Unfortunately, there are no direct measurements for the mobilization parameter (M) that have been obtained independent of HGT. Most studies have the following design in common ^17–21^; the resistance encoded by the donor is not mobile and can only be detected in the recipient once it has been mobilized and transferred. Thus, we do not have a measure of the M parameter, and instead we have used the combined parameter MH (mobilization and horizontal gene transfer) where experimental data can be obtained from these studies.

To get an estimate for the dissemination of ARGs from the environment to humans (D), the number of eaten bacteria per day were used as a proxy ^14^. In this study, the authors counted the number of colony forming units of meals for three different diet types. In this context, other transfer routes are also conceivable, e.g., the absorption of bacteria by swallowing water while bathing ^22,23^. Such dissemination pathways can be of decisive importance when considering individual systems. However, they are not explicitly further considered in this model, which is intentionally kept general, as we aim to accommodate all possible dispersal scenarios in one single parameter.

Finally, some parameters were calculated based on the estimates of the number of humans on earth (7.8 billion; http://www.worldometers.info), the number of bacterial cells in/on the human body (3.8 x 10^13^) ^24^ and the total number of bacterial cells on earth (on the order of 10^30^) ^25^. These values were used to define ranges for Pph, Ph, Pp and E (Table 2). We also estimate a typical bacterial genome to contain 1500 to 7500 genes ^26^, that around 7% of bacterial cells carry a conjugable plasmid, and that a typical plasmid carries 50-100 genes ^27^, which was used to constrain the Pm parameter. The S parameter represents the overall fitness impact of carrying an ARG and implicitly also involves the survival rate for a bacterium carrying an ARG, relative to a non-carrier. S is defined as the population expansion rate per day in the model, so when S is equal to one the average ARG would have no impact on the expansion of the population, i.e. the average ARG would overall be fitness neutral.

### Many resistance genes are likely to have been present in human-associated bacteria at the beginning of the antibiotic era

It is clear from running the model that for many parameters the permitted range of possible values is huge (Fig. 2A). Even for intuitively ridiculous models such as the assumption that all bacteria originally carried all resistance genes (E = 1), there seem to be valid model scenarios. However, it is clear that in those cases, this would have to be compensated by the vast majority of ARGs having a strongly negative effect on fitness, as evidenced by the strong negative correlation between E and S (Fig. 3; Supplementary Fig. 1). As the model results are highly dependent on certain parameters, it is difficult to say what the typical result of the model would be. However, assuming that the dispersal parameter D is in the range between 10^-12^ and 10^-11^, the most likely origins for the currently circulating resistance genes at the start of the antibiotic era would be on mobile genetic elements (MGEs) in human-associated bacteria (65.6%) and on MGEs in environmental bacteria (28.4%; Table 3). These two origins are substantially more likely than ARGs having been present in pathogens at the start of the antibiotic era (3.07%), coming from the chromosomes of non-pathogenic human-associated bacteria (2.03%) or from the chromosomes of environmental bacteria (0.883%). In almost all runs of the model, the likelihood of ARGs originating from pathogenic bacteria in the environment is less than 0.00001% (Table 3). However, the possible ranges for all the other five processes are highly overlapping (Fig. 2B).

**Fig.2.**
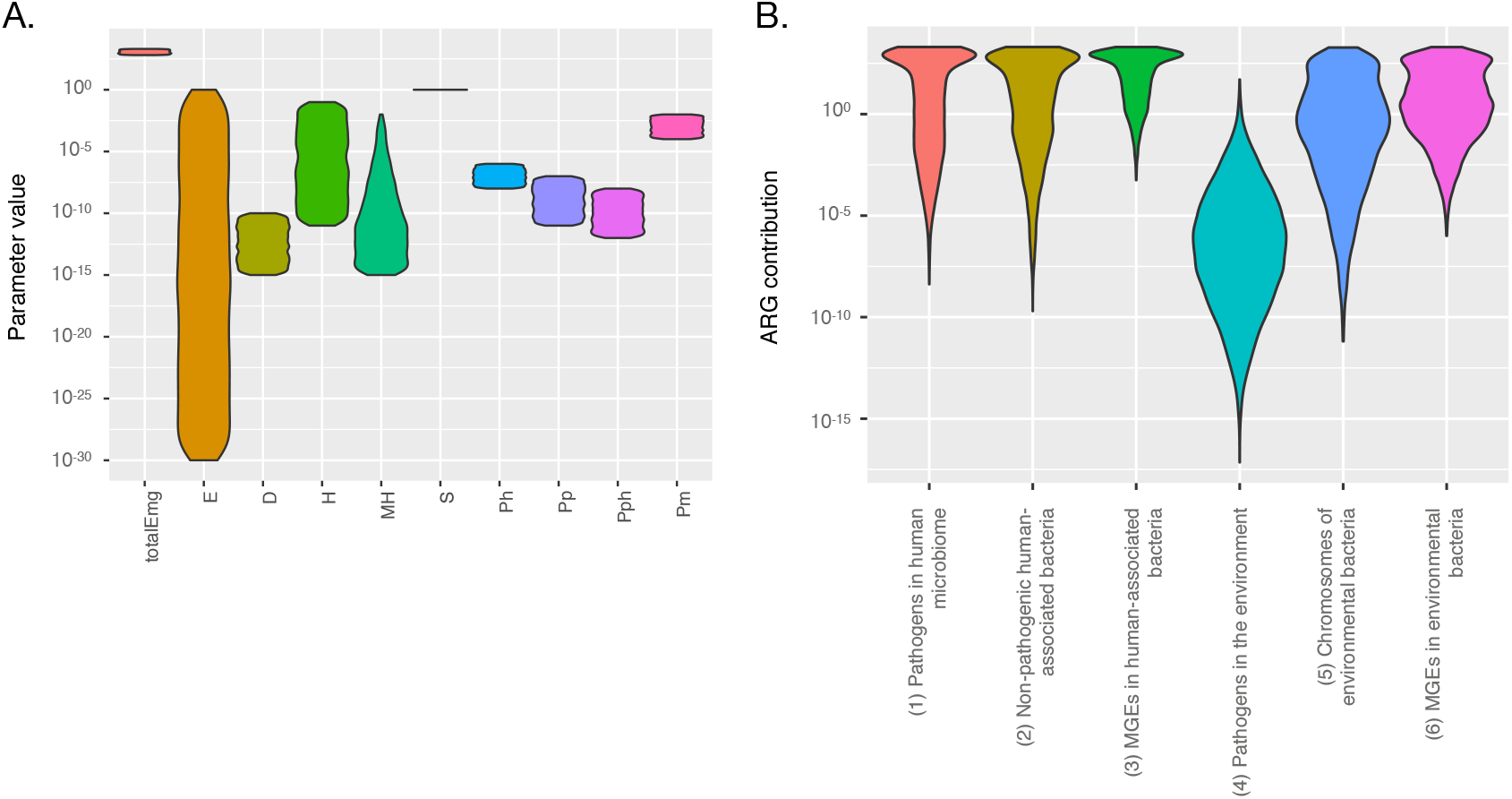
Valid parameter ranges (A) and process rates (B) for the main model after 70 years of simulated time. The range of S extends from around 0.99 to 1.002.

**Fig.3.**
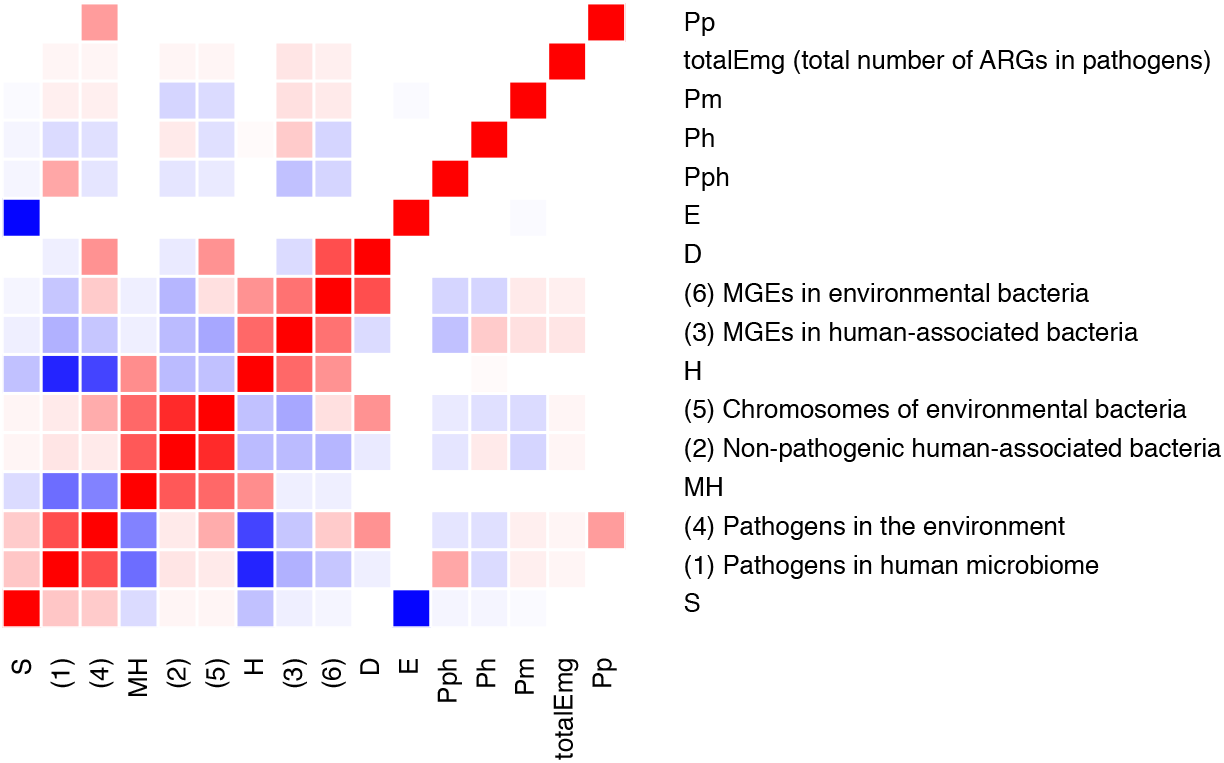
Correlations between the variables in the main model after 70 years of simulated time.

**Table 3.**
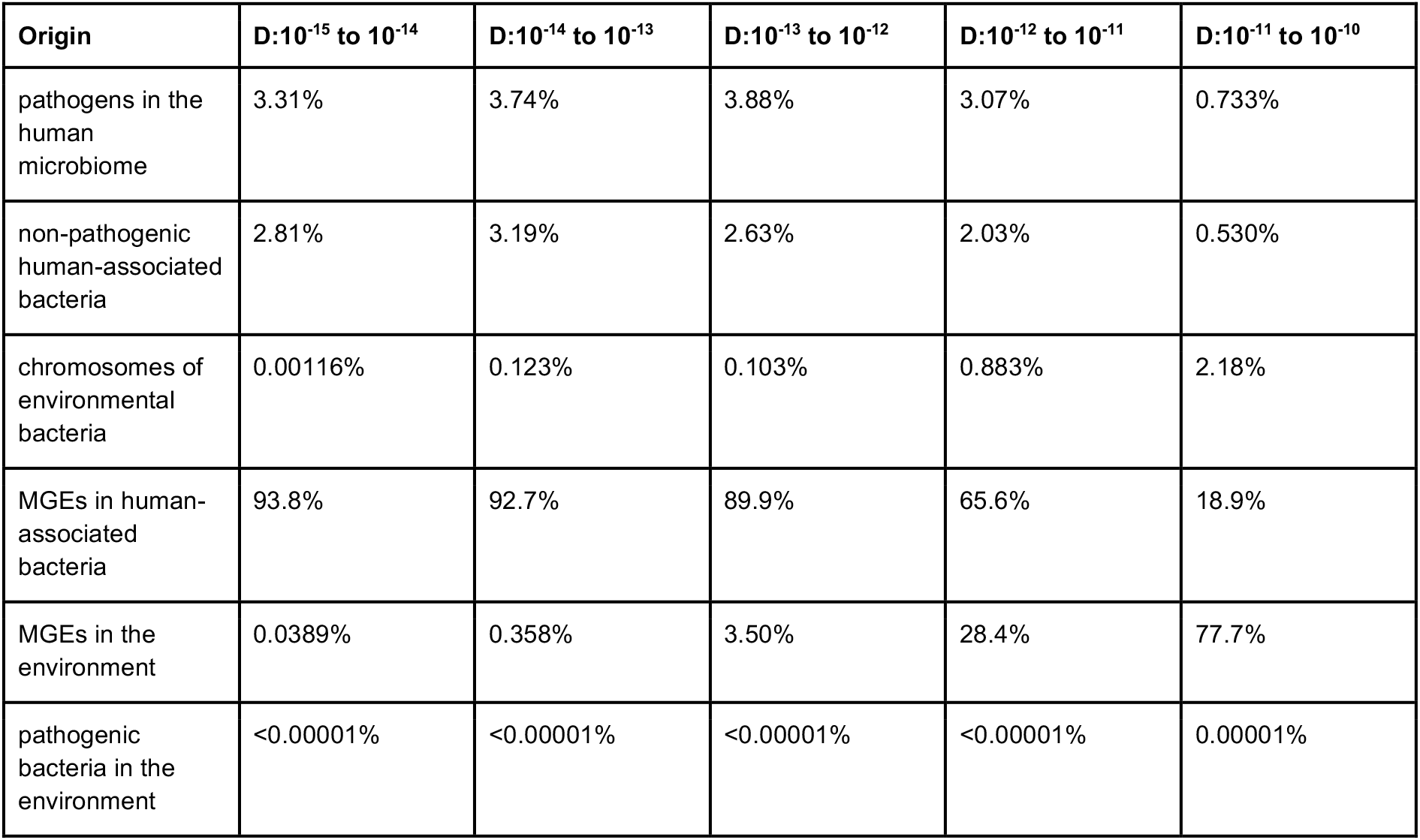
Dependency of the modelled processes on the D parameter

| Origin | D:10<sup>-15</sup> to 10<sup>-14</sup> | D:10<sup>-14</sup> to 10<sup>-13</sup> | D:10<sup>-13</sup> to 10<sup>-12</sup> | D:10<sup>-12</sup> to 10<sup>-11</sup> | D:10<sup>-11</sup> to 10<sup>-10</sup> |
| --- | --- | --- | --- | --- | --- |
| pathogens in the human microbiome | 3.31% | 3.74% | 3.88% | 3.07% | 0.733% |
| non-pathogenic human-associated bacteria | 2.81% | 3.19% | 2.63% | 2.03% | 0.530% |
| chromosomes of environmental bacteria | 0.00116% | 0.123% | 0.103% | 0.883% | 2.18% |
| MGEs in human-associated bacteria | 93.8% | 92.7% | 89.9% | 65.6% | 18.9% |
| MGEs in the environment | 0.0389% | 0.358% | 3.50% | 28.4% | 77.7% |
| pathogenic bacteria in the environment | <0.00001% | <0.00001% | <0.00001% | <0.00001% | 0.00001% |

Furthermore, the interdependencies between the process rates and some of the individual parameters are, in many cases, fairly high. For example, the likelihood of an ARG originating from the chromosomes of environmental bacteria or from non-pathogenic human-associated bacteria is strongly correlated to the value of MH – the mobilization rate of genes to MGEs, including subsequent gene transfer (Fig. 3). Similarly, the proportion of ARGs that already existed in pathogens in humans to begin with is, unsurprisingly, dependent on the proportion of all bacteria that are human pathogens (Pph), and the likelihood of originating on an MGE – either in a human-associated bacterium or in the environment – is strongly dependent on the rate of horizontal gene transfer (H). Importantly, though, the number of bacteria carrying a certain ARG from the start of the modelling time (E) is almost entirely dependent on the fitness cost or advantage associated with the gene (S), highlighting that selection for ARGs is by far the most important process in ARG ecology. The parameter that largely controls whether the environment is likely to be a significant contributor to ARGs in pathogens is D (Fig. 3). If the environment played a major role in the emergence of the nowadays known ARGs in human pathogens, dissemination from the environment to humans (D) as well as the mobilization and transfer rates (MH) of ARGs need to be fairly high. It is also notable that the three parameters that influence the outcomes of the model the most are S, H and MH (and indirectly E, as it is anti-correlated with S; Fig. 3). In particular, changing the values of MH and H shifts the relative importance of the processes and thus the most likely origin of ARGs dramatically (Supplementary Table 1). Again, it should be pointed out that the span of possible outcomes for a given size of MH, H or D is still very large (Supplementary Figs. 2–4).

### Stability of the parameters to changes

We also investigated how stable the model was over time (i.e. when the number of days were varied in the model). Again, the strong relationship between the E and S parameters was evident (Fig. 4A and B). Particularly, the value of S becomes highly constrained over long timescales, hinting at that most ARGs that have made it all the way to human pathogens should be close to neutral with respect to impact on host fitness. It is also interesting to note the relative increase in environmental processes over time (Fig. 4C), which also relates to the impact of selection over time and how the environment presents a vastly larger number of bacteria with potentially neutral ARGs to recruit from.

**Fig. 4.**
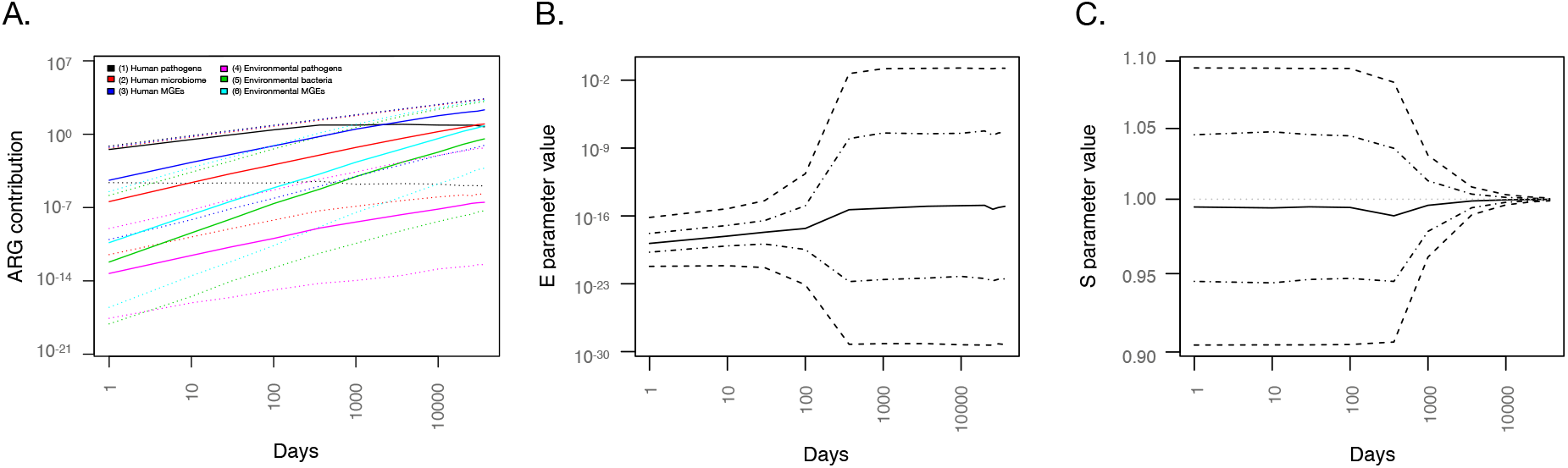
Dependence of model process rates (A) and key model parameters E (B) and S (C) over the simulated time span from the start of antibiotic use to the emergence of ARGs in human pathogens. Dotted lines in (A) and dashed lines in (B) and (C) represent the range in which 95% of the values fall in the simulations. Lines with dots and dashes in (B) and (C) represent the range in which 50% of the values fall in the simulations.

Building on the assumption of neutral impact on host fitness (or, more precisely, neutral *average* impact on fitness over the run time of the model), we also ran the model with S fixed to 1 (i.e. no effects on fitness for any ARG). This is an interesting scenario as it caps E to the range between 10^-24^ and 10^-11^ (Supplementary Fig. 6), meaning that if fitness cost is neutral, currently observed ARGs in pathogens would have been present in somewhere between 10^6^ and 10^19^ bacterial cells at the start of the antibiotic era. However, other than that fixing S to 1 changes how the different processes depend on the E parameter (Supplementary Fig. 7), the model output does not substantially change compared to running the model with a variable S parameter.

We also compared the main model to the emergence model, in which ARGs are assumed not to have been already present in bacteria at the start of the antibiotic era, but instead had to develop their resistance function – emerge as ARGs – within that time frame. However, the results differed only slightly from the results of the pre-existing model (Supplementary Fig. 8). Based on the model results, the conceptual difference between the pre-existing model and emergence model is not sufficient to change the dependencies between the rest of the processes. The full results from the emergence model are available in the supplementary material.

### Model predictions

Notably, the model makes some predictions that may be testable within the foreseeable future. The respective results will substantially contribute to determine the crucial processes in antibiotic resistance development and thus help in the identification of effective intervention strategies:

1. Fitness costs of ARGs: The model predicts that the majority of ARGs present in pathogens today should have very limited effects on fitness. The model caps the average fitness impact for ARGs currently present in human pathogens between −0.2% and +0.1% per generation, and 30 years into the future the model predicts that the range of possible fitness effects will have narrowed even further (−0.15% to +0.1%; data not shown). By determining the fitness effects of carrying individual ARGs in their current hosts considering inter-as well as intra-species variability, the prediction that most ARGs impose a very minor fitness impact could be experimentally tested, although it would have to be tested across a large number of ARGs.
2. The origin of ARGs: The model predicts the most likely location of ARGs 70 years ago would have been in human-associated bacteria (Table 3). By tracking ARGs currently present in human pathogens across large datasets of bacterial genomes it may be possible to trace the evolutionary history of these genes and thereby identify their likely hosts at the beginning of the antibiotic era. Such an attempt was recently made, corroborating the findings based on our model ^28^ and lending support to that most ARGs may not originate in the environment. However, this analysis is complicated by the biased sampling of fully sequenced bacterial genomes, most of which originate from human specimens (https://gold.jgi.doe.gov/distribution). Thus, it could be expected that when performed today, such an analysis would only be able to trace origins for ARGs that were present in human-associated bacteria 70 years ago. Notably, the model estimates that such genes would be more likely than not to make up the majority of all ARGs currently present in pathogens. Importantly, the rapid increase in sequencing capacity may make full-scale analysis of ARG origins using genomic data possible in the relatively near future, which would enable further testing of this prediction of the model.
3. More data will enable additional specific predictions: Given that the origins of ARGs currently circulating in pathogens can be established, the range of possible predicted values of D narrows dramatically. If it can be shown that the proportion of ARGs in pathogens today that were already present in human-associated bacteria 70 years ago is not even close to 60%, the dispersal parameter D has to be above 10^-12^ (Table 3). Conversely, if M and H could be better determined by experiments, the model would predict the likely origins more precisely (Supplementary Table 1). Just establishing a ball-park range of the mobilization rate (M) would dramatically restrict the possible outcomes of the model, particularly if M turns out to be at the extreme ends of the spectrum. Thus, a more precise determination of any of these parameters would enable several more specific predictions by the model,

## Discussion

In this paper, we present a model for ARG appearance in pathogens, based on the limited data currently available on process rates in the literature. The model predicts a vast range of possible outcomes and parameter values; however, it serves its purpose to highlight which processes and parameters that are interdependent and which ones are key to get a better understanding of how to mitigate antibiotic resistance development. A key aspect shaping the model is the strong relation between longevity of ARGs (intrinsically largely determined by their fitness cost) and how widespread resistance genes were across bacteria at the start of the antibiotic era (which likely serves as a proxy for how large the total genetic reservoir to recruit ARGs from still is today ^29^). As the average fitness cost of carrying an ARG increases in the model (resulting in lower values of the S parameter), the number of available ARGs – either in the human microbiome, the environment, or both – needs to get bigger to compensate for that. As the model scenario plays out over a vast amount of time (in microbial terms), the possible range of average fitness costs or gains associated with ARG carriage becomes very small. Thus, the model essentially predicts that most ARGs present in pathogens today should have no or little fitness cost, which is in accordance with theoretical arguments we have put forth previously ^9,11^. As mentioned earlier, this prediction is, in principle, testable by large-scale experiments on the impacts of ARG carriage on bacterial fitness. Notably, low average fitness costs for ARGs are associated with a greater likelihood that they have been present in human pathogens all along. Overall, it is clear that the fitness cost of ARGs is a key property for their ability to make their way to and persist in human pathogens. The persistence is likely to be facilitated by exposure to low levels of antibiotics, but also by mutational processes that lead to domestication of ARGs and MGEs, as well as other forms of fitness cost reductions, some of which may yet be unknown. In this context, a variety of mechanisms to reduce the fitness cost of ARGs are known, but quantitative information on their occurrence and effect is still largely lacking (Table 4). More research is needed to quantitatively determine the impact of domestication and reduction of fitness costs on selection of ARGs, as well as what additional mechanisms that are available to bacteria for reducing fitness costs of ARGs. Furthermore, minimal selective concentrations must be determined for a much larger set of antibiotics, environmental conditions and bacterial species, alone as well as in microbial communities, to provide actual scientific data in a field that currently relies largely on predictions of antibiotic effects on microbes ^30,31^.

**Table 4.** Identified knowledge gaps and research needs that must be addressed to build quantitative risk assessment models and better mitigate ARG recruitment

| Knowledge gap | Research need | Utility |
| --- | --- | --- |
| Settings that select for antibiotic resistance | Minimal selective concentrations (MSCs) for more antibiotics, environmental conditions and bacterial species, alone, in co-culture and in complete communities | Better sewage and wastewater treatment strategies; emission limits and environmental quality standards for antibiotics |
| Rate of bacterial dispersal between environmental compartments (and to humans) | Quantitative observations of bacterial dispersal between environments; tracing spread of ARGs between environments over time; better determination of human exposure to environmental bacteria | Ability to discern if the environment has a significant role in the emergence of novel ARGs; improved quantitative risk assessment models for environmental antibiotic resistance; ability to limit spread of ARGs and resistant bacteria from the environment to humans; improved wastewater treatment strategies |
| Quantitative information on the occurrence and effect of mechanisms for fitness cost reduction and domestication of ARGs | Quantitative understanding of genomic mechanisms and genes responsible for fitness cost reduction of ARGs | Improved quantitative risk assessment models for environmental antibiotic resistance |
| Rate of horizontal gene transfer between bacteria | Measurement of transfer rates (especially for transformation and transduction) between a larger diversity of bacterial species and in a greater number of settings | Improved quantitative risk assessment models for environmental antibiotic resistance; ability to curb transfer of ARGs between bacteria or reduce the number of settings where this can take place |
| Rate of mobilization of genes from bacterial chromosomes to MGEs | Development of proper experimental setups to measure mobilization; measurement of mobilization rates in such assays | Improved quantitative risk assessment models for environmental antibiotic resistance; better understanding of whether the environment acts as a source of ARGs to human pathogens, and thereby better prioritization of interventions |

Along with the importance of fitness costs and selective effects, the model also highlights the fundamentally important impact of the dispersal parameter (D) on ARG occurrence in pathogens. This parameter essentially determines if the environment would at all plausibly function as a reservoir for ARGs to be recruited to pathogens, as has been argued by many authors ^4,32–34^. Notably, there are more scenarios in the model where the vast majority of ARGs originated in human-associated bacteria than there are scenarios where the environment played a significant role, although both situations are possible. Interestingly, a recent study using genomic information to infer the recent origins of ARGs assigned more than 90% to bacteria associated with humans and/or domestic animals ^28^, which is consistent with the most common results of our model. While there are some limitations to the currently available genomic data, this implies that the environment would only play a minor role in the development of new ARGs and, consequently, that the D parameter is likely to be smaller than 10^-12^.

Importantly, the data on dispersal of genes from environmental bacteria to the human population is very limited, particularly for the type of generalized case that we outline in this paper. Due to the lack of data, we have chosen a fairly naive definition of dispersal for our model, reflecting the type of data that is available. The D parameter in the model represents the proportion of environmental bacteria that would be expected to be exposed to a human somewhere in the world during a day. This was estimated from the number of cultivable bacteria contained in different types of food as reported by Lang et al.^14^. However, depending on the diet investigated, the approximate numbers of bacteria taken in from food were in the range of 10^6^ to 10^9^ bacteria per person and day, which is a fairly wide range. In addition, this does not account for bacterial exposure through other routes, such as recreational swimming ^22,23^, wounds ^35^ and through direct contact with animals ^33,36,37^. Thus, the definition of D in our model does not reflect that the origins of bacteria that the average human is exposed to are not equally distributed across environments. To some extent, we expect this to be compensated by bacterial global dispersal over time, but this still reflects a weakness in our model. That said, although it is feasible that all these activities contribute significant numbers of bacteria, it is unlikely that they would change D by several orders of magnitude. However, for this reason it may be fair to assume that D is in the higher end of the spectrum, i.e. larger than 10^-12^ rather than close to the lower boundary. If D would be larger than 10^-11^, environmental bacteria may have contributed around 80% of all currently clinically relevant resistance genes (Table 3). This, however, runs contrary to recent genomic evidence ^28^, although the latter is based on a likely biased sample of genomic data. The impact of the D parameter highlights that determining the actual dispersal rate of ARGs from the environment is crucial to understand the role of the environment in antibiotic resistance development. Comprehensive data on ARG dispersal would not only aid in risk assessment of emergence of novel resistance factors; it would also aid in understanding and mitigating the spread of resistant bacteria (and their ARGs) through the environment ^6,8,10,38^.

The model also reveals that HGT and mobilization of chromosomal ARGs to MGEs are potentially important processes, although likely less to be bottlenecks in resistance development than selection and dispersal. While we have a hunch about what the HGT rates may be (ranging from 2.4*10^-10^ to 5.8*10^-2^ events per cell per day in our literature review; Supplementary Table S2), this range is still not sufficiently narrow to really restrict the model outcomes in a meaningful way. That said, at high rates of HGT, most ARGs are, according to the model, likely to have originated on MGEs, unless the mobilization rate is also high. Unfortunately, the latter parameter is almost completely unknown – the little available data we could find suggest that the rate of mobilization followed by HGT is in the range between 4*10^-10^ and 1.5*10^-4^. Since these estimates are all based on mobilization followed by HGT, they do not represent the mobilization rate *per se*, and thus the actual rate may turn out to be outside of the mentioned range. In addition, these rates have been measured using very specific markers that are known to be transferable to plasmids, which may not make them representative for the mobilization rate of ARGs in general. Clearly, there is a need to better define the typical rates of HGT between bacteria. Several assays already exist to perform such experiments ^39–42^, but the amount of data available is still insufficient as it is only available for a very limited number of conditions, ARGs and mating pairs. Furthermore, most research on HGT has dealt with conjugation, while transformation and transduction has been largely neglected, particularly from a quantitative point of view. Still, the two latter processes have been shown to be important drivers of the horizontal transfer of resistance genes ^16,43–45^, although their contribution relative to conjugation is unclear. Importantly, this is largely a data generation problem. In contrast, the lack of knowledge of the rate of mobilization of chromosomal genes to MGEs is not only related to insufficient data, but also to a lack of appropriate assays to actually measure mobilization of ARGs rather than mobilization linked to, e.g., horizontal gene transfer. This highlights an urgent need for assays to directly detect mobilization of genes from the chromosomes to MGEs.

It is important to note that we have intentionally kept the model and its parameters general, in order to provide an overview of the respective importance of the individual processes that influence the appearance of new resistance genes in human pathogens. This is likely also one of the reasons why the model predictions are in many cases associated with very wide ranges. However, the model still emphasizes important and interrelated processes that need more research attention in order to build more quantitatively accurate models. Furthermore, by using specific rates for particular ecosystems as input for the model, we believe that our model could be used to assess the probabilities for the contribution of specific pathways for resistance development and dissemination. Importantly, this model constitutes an initial effort to create a conceptual framework for ARG emergence in pathogens ^9,11^ that is quantitative, and to highlight where more data is needed in order to make more precise predictions. As such, we hope that this modelling effort can spur more research; both into determining the model parameter ranges more precisely, but also into more sophisticated modelling approaches that may better reveal the details on ARG emergence.

Another limitation to the model is that the variable associated with the fitness cost of ARGs (S) represents a variety of different processes and parameters related to bacterial survival and proliferation. As we did not find an unambiguous way to tease these components apart, they are all captured by this single parameter, which of course results in an oversimplification of the real world. For these reasons, the role of S has to be interpreted with some caution. First of all, S represents the *average* fitness impact of the *average* ARG over the entire run time of the model (which in most of the data we have presented in this paper is 70 years). This means that S encapsulates scenarios where an ARG may some of the time have a positive fitness impact (such as during antibiotic selection) and at other times a slightly negative effect on fitness (e.g., in the absence of antibiotic selection). The final value of S therefore represents the expected overall fitness impact of ARG carriage, not the specific fitness of any single ARG currently encountered in pathogens. Furthermore, S also implicitly includes a constraint on the survival rate for a bacterium carrying an ARG. The way S is defined in the model, it is related to the population expansion rate per generation. In other words, if S is equal to one, the average ARG is expected to have no impact on bacterial growth and, thus, the population expansion (i.e. it is fitness neutral). However, given the relatively quick doubling rate of bacteria, a value only slightly deviating from one will rapidly result in the gene propagating through bacterial populations or quickly disappearing, which explains the tendency of S to approach one the longer the simulated time of the model.

Despite the large uncertainties associated with the model results, some useful intervention priorities can still be derived from this work (Table 4). Firstly, due to the strong influence of selection and ARG fitness costs on the model outcomes, it is clear that there is a need for a deeper understanding of the settings that drive antibiotic resistance selection. Identifying threshold concentrations of antibiotics (and other selective agents, such as biocides ^27,46^) that select or co-select for antibiotic resistance is crucial for determining which emission limits that should be imposed on effluents from pharmaceutical production ^47,48^ as well as from regular wastewater treatment ^49–51^. Currently, regulatory efforts are mostly based on predicted no-effect concentrations ^30,31^, and while this is a decent interim solution, solid scientific data on actual minimal selective concentrations in relevant settings are urgently needed ^10^.

Filling the knowledge gaps in terms of human exposure to environmental bacteria carrying ARGs also has important implications for prioritization of interventions to prevent the emergence of antibiotic resistance in pathogens. The data we present in this paper suggest that a low dispersal rate renders the environment implausible as a source for the vast majority of ARGs in pathogens. If this is true, that would imply that in the short term, interventions targeting human use of antibiotics should be prioritized over, e.g., environmental interventions, as the latter might be ineffective in preventing resistance emergence in pathogens. That said, the environment may still function as a dispersal route for already resistant pathogens, and thus environmental mitigations should be selectively targeted towards minimizing spread of resistant bacteria from the human population to the environment.

The model furthermore implies that a low mobilization rate would cause the spread of ARGs from resistant bacteria to susceptible pathogens to be the most important vehicle for recruitment of novel ARGs. This may also be the case if mobilization rates are high, but then to a lesser extent. These observations highlight the importance of improving our determination of the typical range of HGT rates of ARGs between bacteria. The impact of environmental conditions and stressors on the transfer rate is also crucial to determine, as that knowledge would potentially allow interventions to reduce the number of settings where transfer between bacteria can take place, and thus delay resistance emergence in pathogens.

Finally, if we can gain a better understanding of how resistant bacteria and ARGs disseminate via the environment between compartments and, particularly, from the environment to humans, it would be possible to start designing mitigation strategies aiming to reduce the spread of resistant bacteria through the environmental route. Such interventions could be made either on the level of releases into the environment or attempting to create barriers between humans and particular risk settings ^8^. Which of those that would be most beneficial is to a large extent determined by the rate of dispersal of resistant bacteria from the environments to humans. Sadly, our current knowledge of the environmental dissemination routes is poor, mostly anecdotal and virtually without quantitative measurements, which is also reflected in the simplistic definition of the dispersal parameter in our model.

## Conclusions and outlook

In this paper, we present a model to describe the emergence of ARGs in human pathogens, and populate it using the limited literature data that exist. The model indicates that fitness costs of ARG carriage is one of the key factors determining the fate of ARGs in both the environment and in human pathogens. It also points to the importance of, particularly, the degree of dispersal of resistant bacteria from the environment to humans, but also the rates of mobilization and horizontal transfer of ARGs. All of these processes are poorly described at present; particularly the possible range of mobilization rates is practically unknown and estimates of dispersal of bacteria from the environment to humans are uncertain. Furthermore, depending on how large these rates are, the model predicts very different probabilities for the possible origins of the ARGs we currently see in pathogens, indicating yet again the importance of determining these values with better precision. However, despite a large degree of uncertainty of specific model results, the model clearly highlights where the most urgent needs for quantitative data exist. The knowledge that could be gained from more precise quantitative process data would be extremely helpful in deciding which processes that should be targeted for interventions of environmental antibiotic resistance development and transmission. By measuring process rates for mobilization, transfer and dissemination, as well as the fitness costs associated with ARG carriage, the model used here could be refined to make more precise predictions on the importance of certain parameters and environments. This will allow for better monitoring of antibiotic resistance and interventions towards preventing it in the future.

## Acknowledgements

JBP acknowledges funding from the Centre for Antibiotic Resistance Research at the University of Gothenburg, the Swedish Research Council for Environment, Agricultural Sciences and Spatial Planning (FORMAS; grant 2016-00768), the Swedish Research Council (VR; grant 2019-00299) under the frame of JPI AMR (EMBARK; JPIAMR2019-109), the Sahlgrenska Academy at the University of Gothenburg, and the Swedish Cancer and Allergy fund (Cancer-och Allergifonden). SH acknowledges funding from the German Research Foundation (HE 8047/3-1)

## Additional figures

**Fig.S1.**
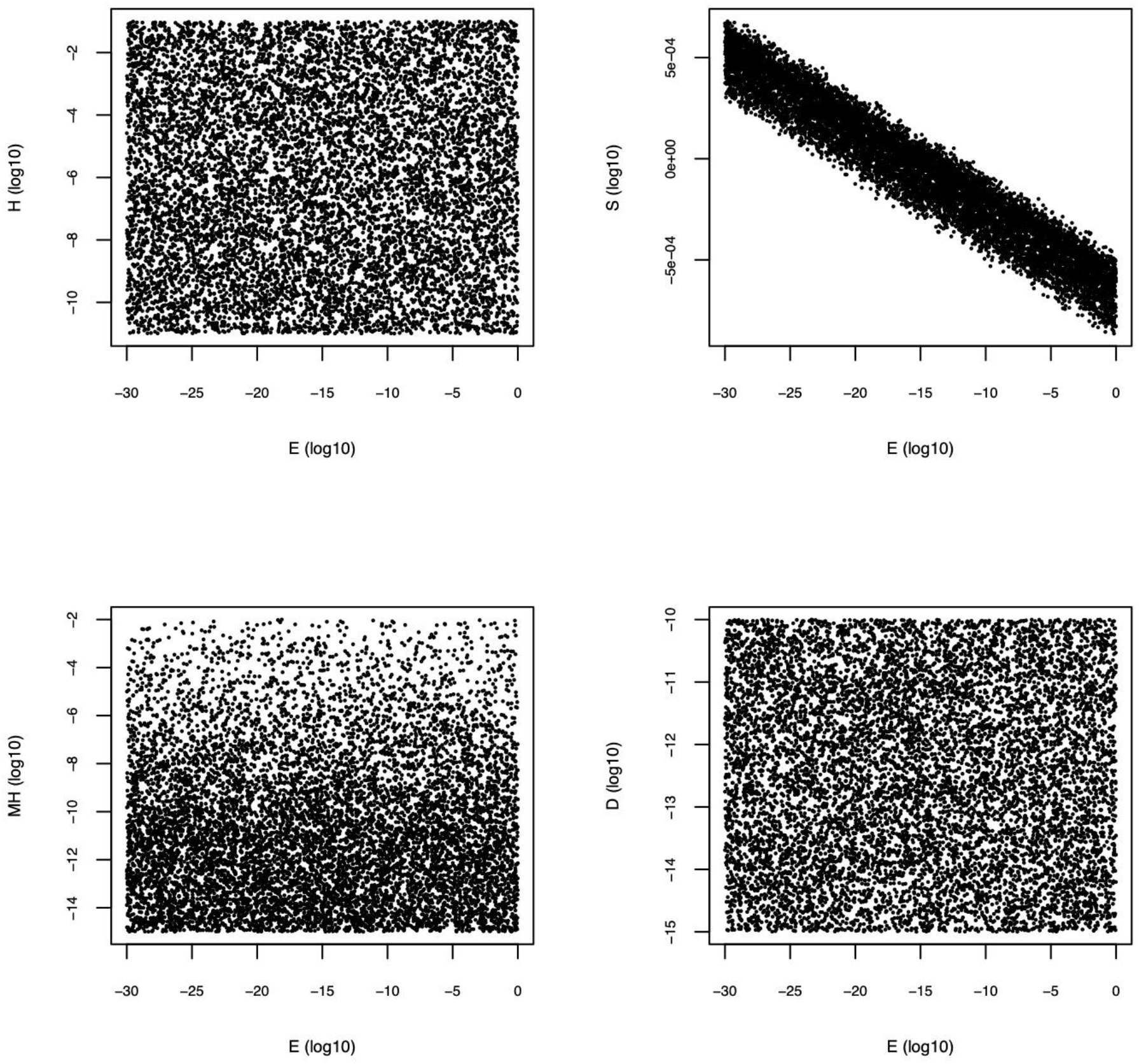
Relations between E and other parameters for the main model after 70 years of simulated time.

**Fig.S2.**
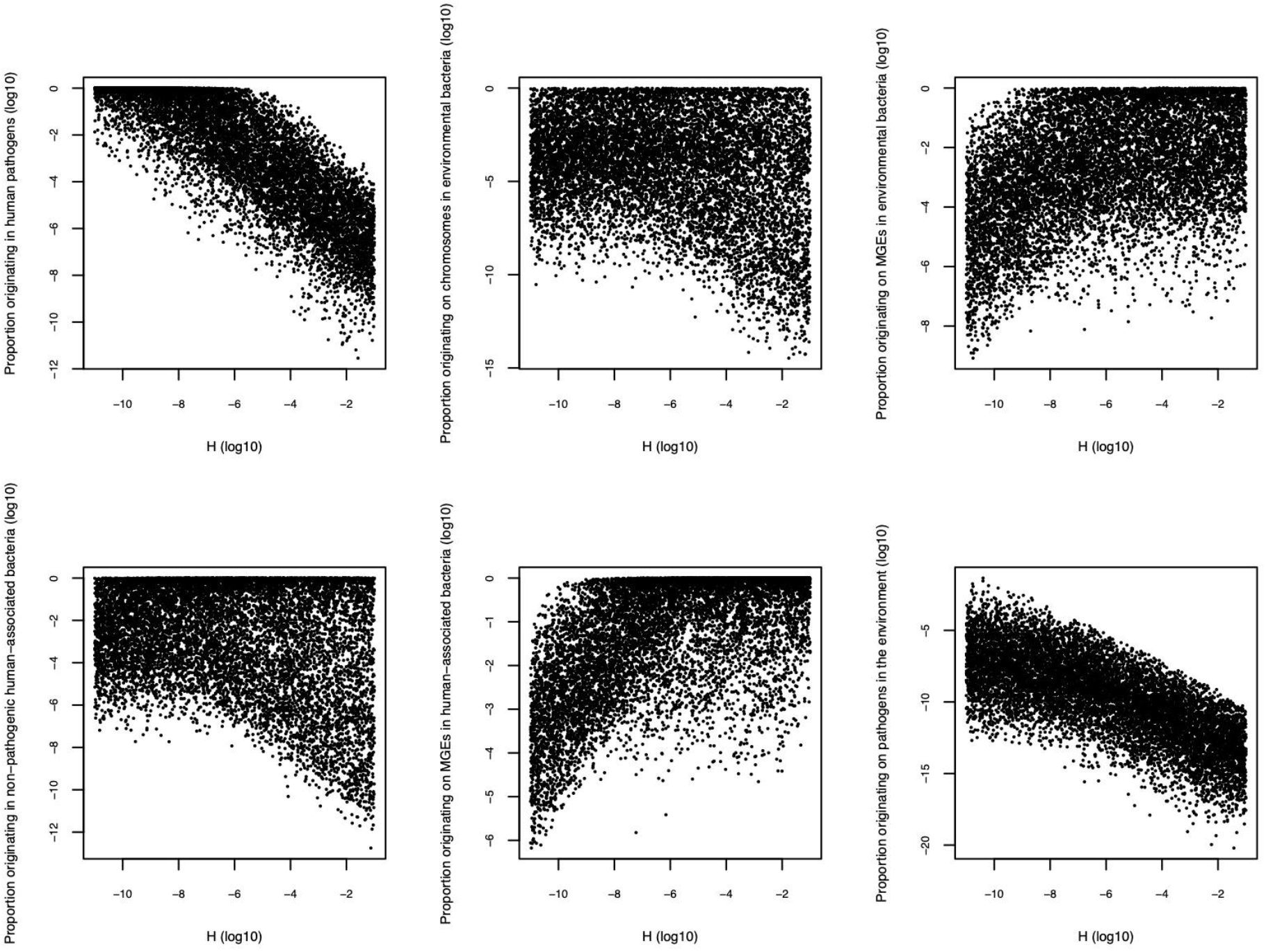
Dependency of the different processes on the H parameter for the main model after 70 years of simulated time.

**Fig.S3.**
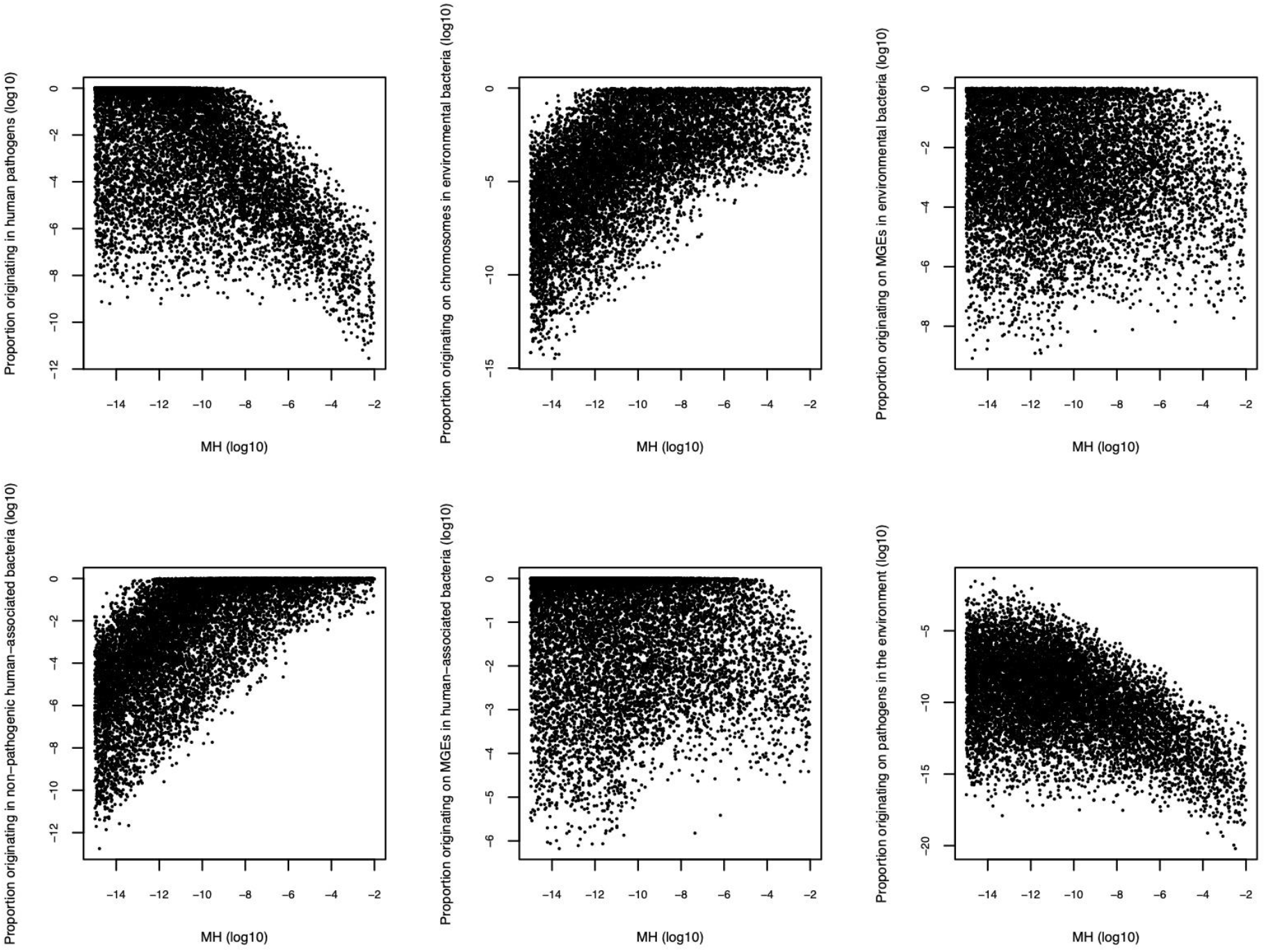
Dependency of the different processes on the MH parameter for the main model after 70 years of simulated time.

**Fig.S4.**
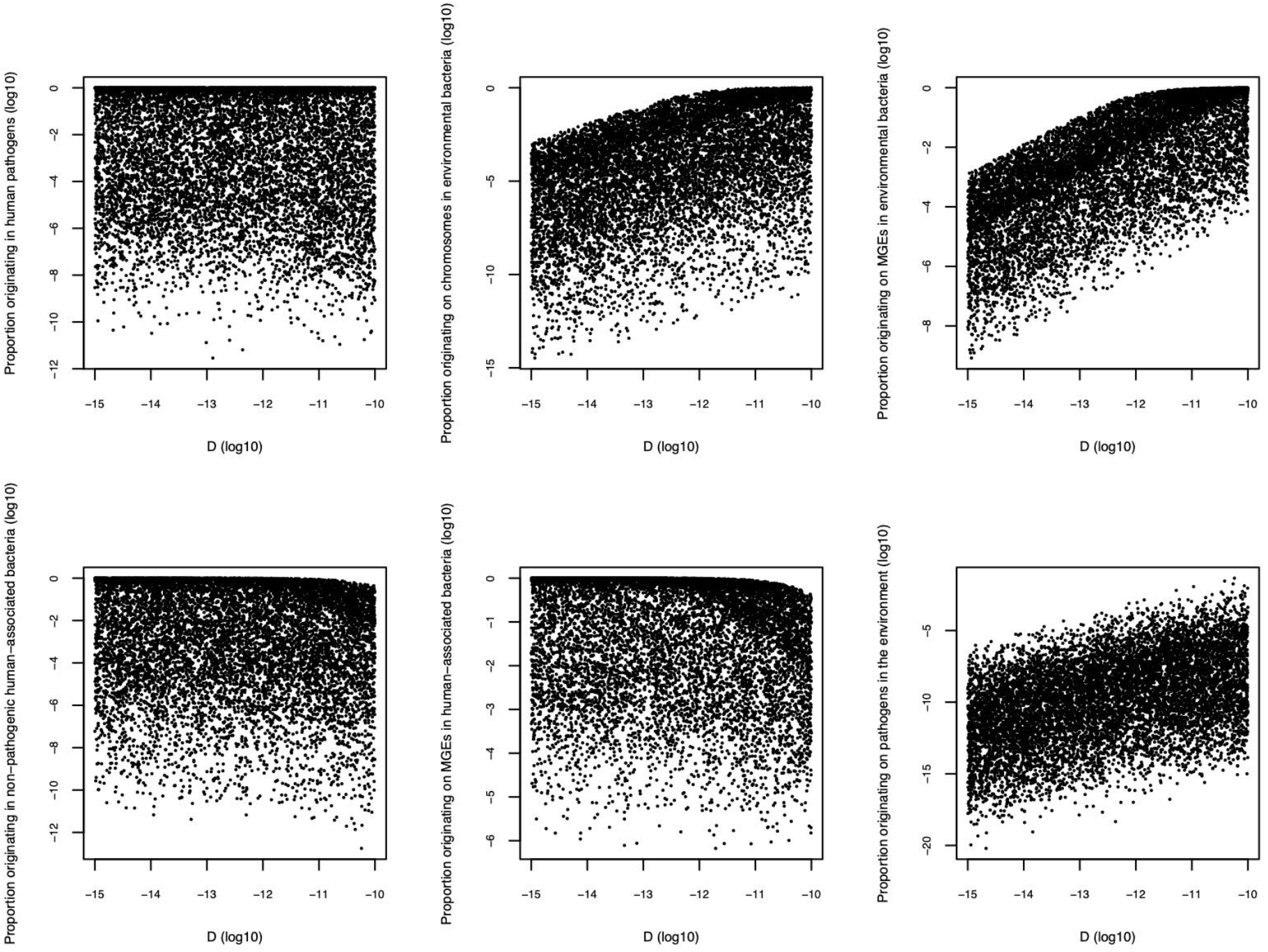
Dependency of the different processes on the D parameter for the main model after 70 years of simulated time.

**Fig.S5.**
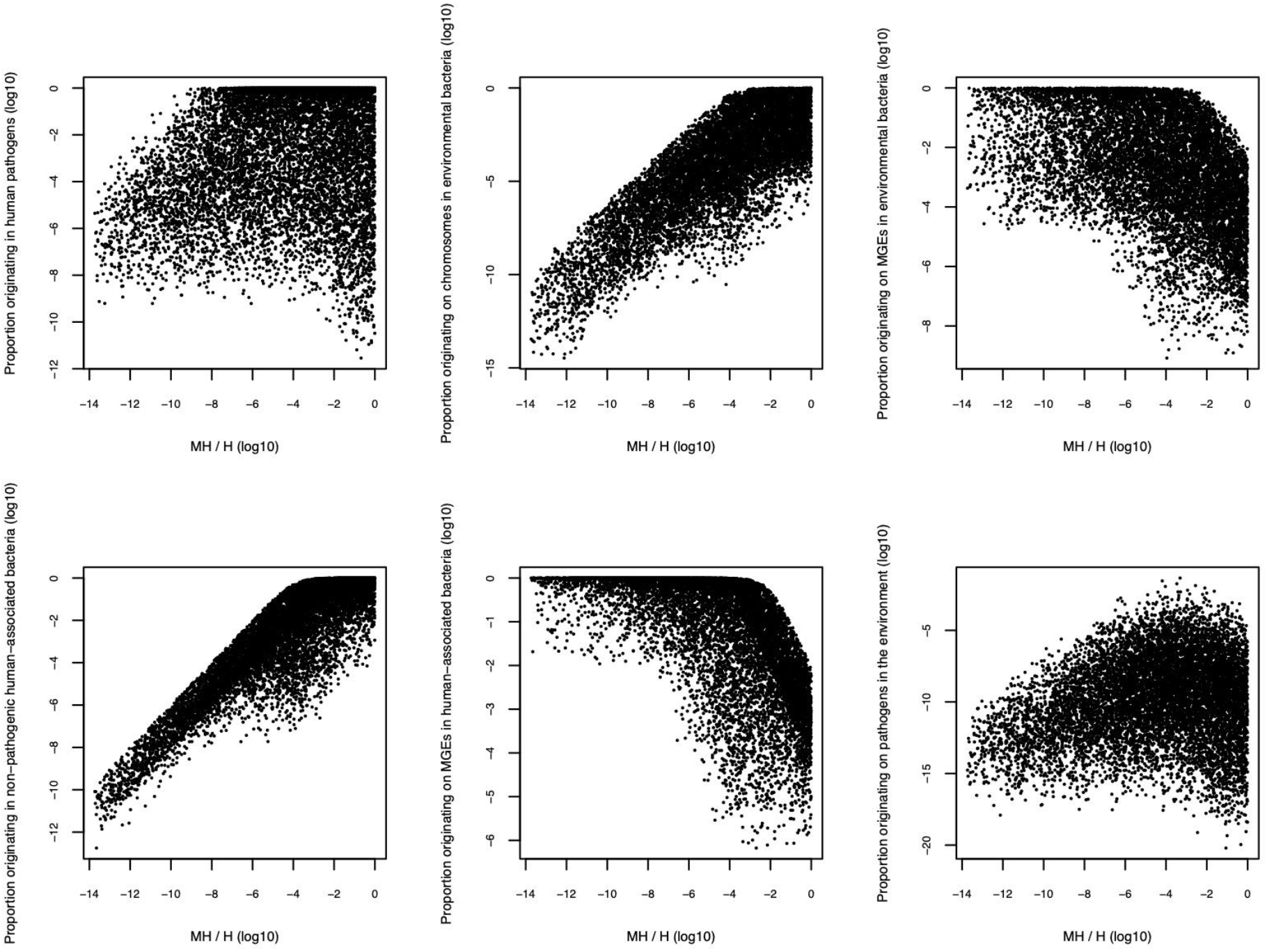
Dependency of the different processes on the MH parameter divided by the H parameter (resulting in an estimated M parameter) for the main model after 70 years of simulated time.

**Fig.S6.**
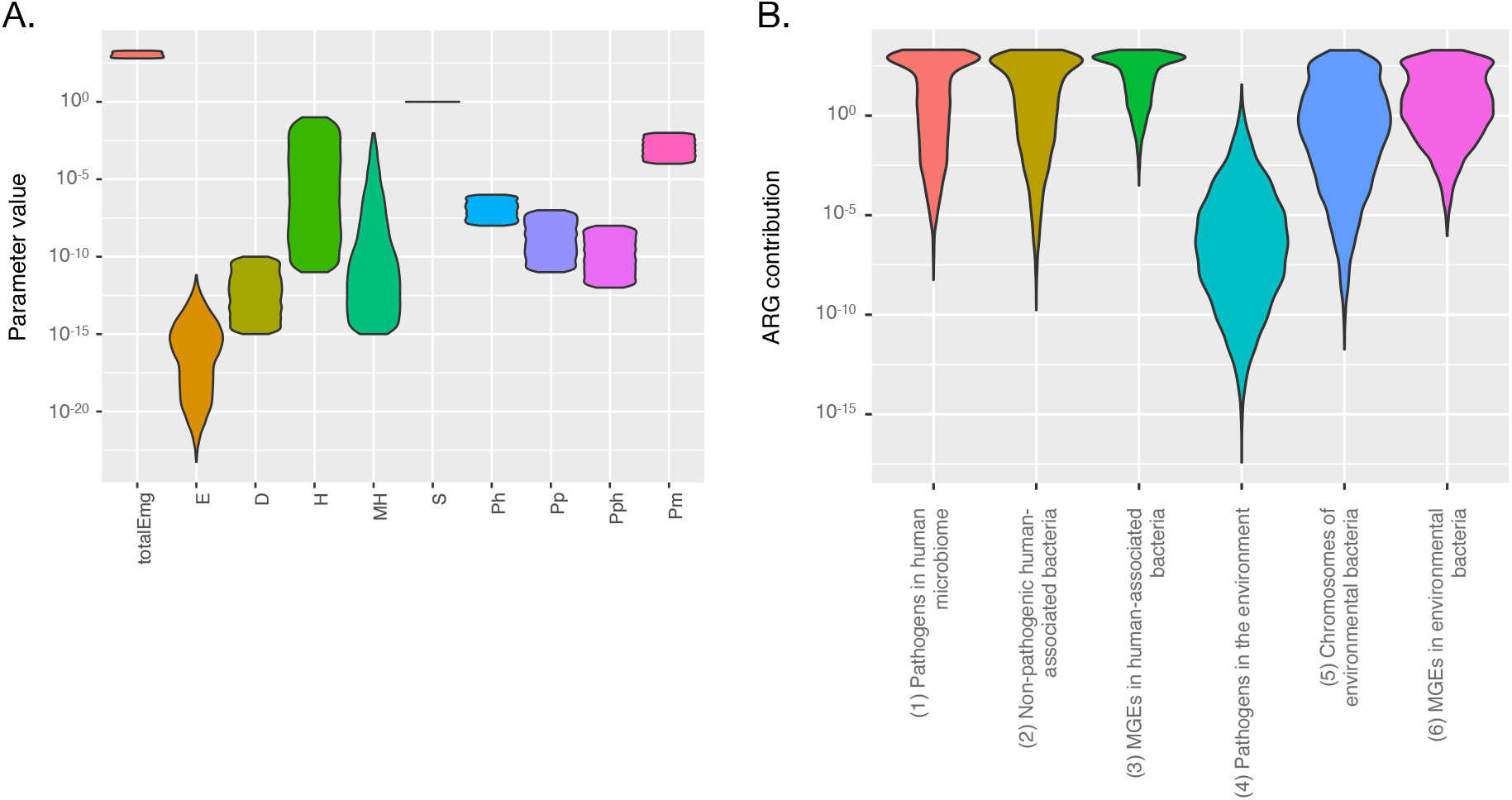
Valid parameter ranges (A) and process rates (B) for the main model after 70 years of simulated time with S being fixed to 1.

**Fig.S7.**
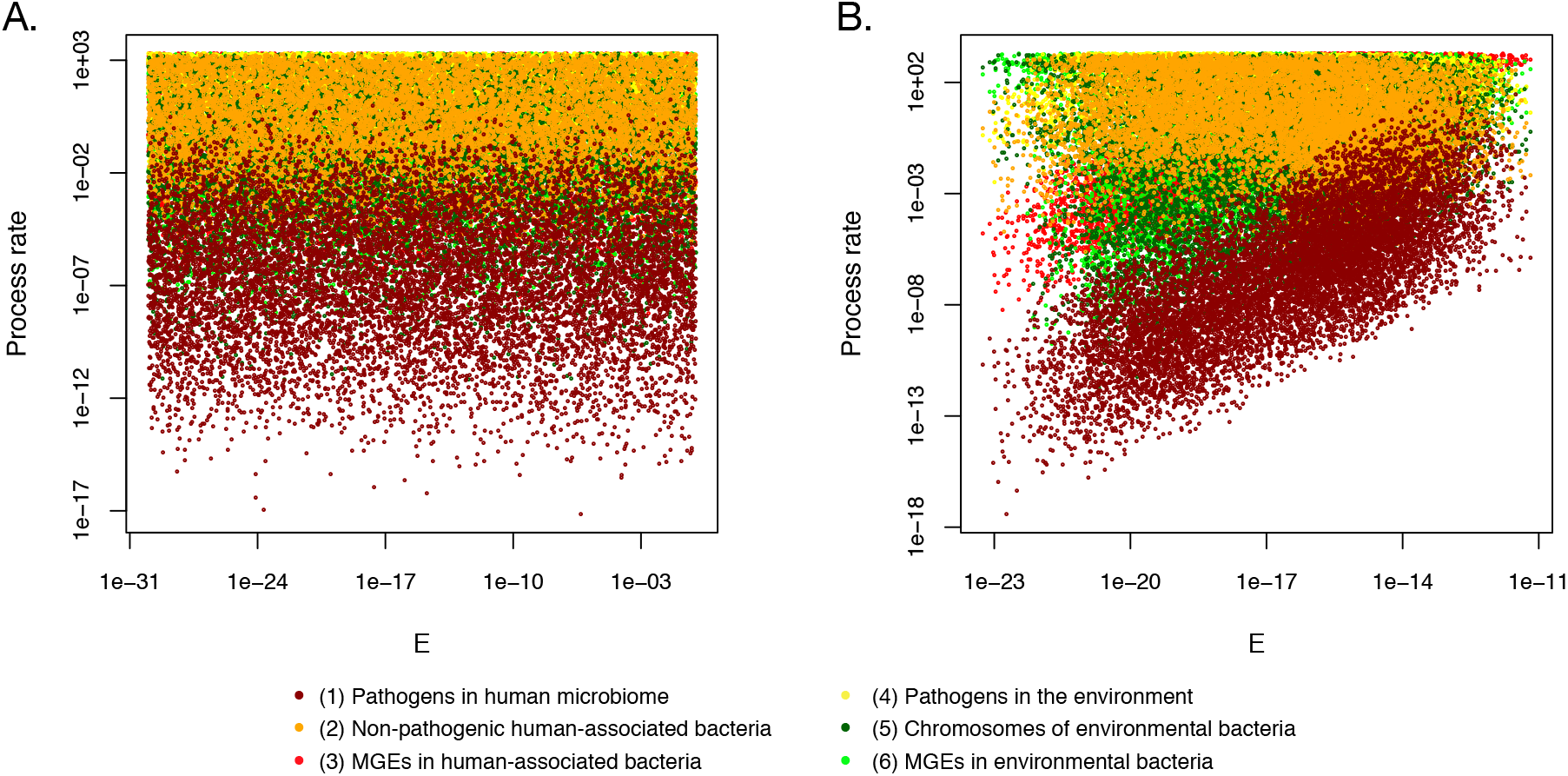
Differences between how processes depend on E for the main model (A), and the model where S is fixed to 1 (i.e. neutral fitness costs of ARGs, B). In both cases, data is presented at 70 years of simulated time.

**Fig.S8.**
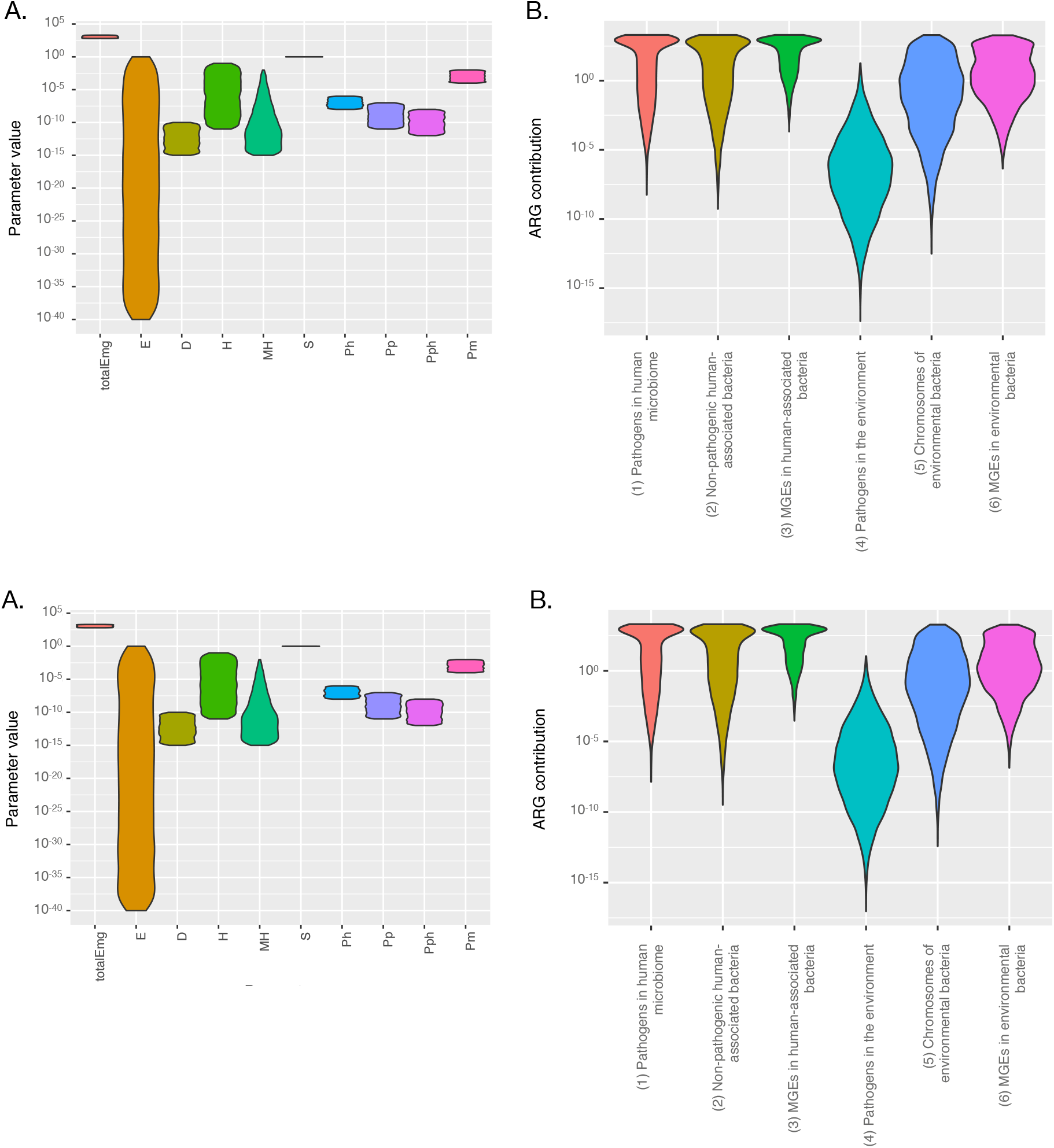
Valid parameter ranges (A) and process rates (B) for the emergence model after 70 years of simulated time.

**Table S1.** Literature survey: Mobilization.

| Study | Species | Test system | Events/Day (average) | Remark |
| --- | --- | --- | --- | --- |
| Torres et al., 1991 <sup>19</sup> | *Enterococcus faecalis* | Tn925 | 1.89E-07 | Also includes a HGT step |
| Manson et al., 2010 <sup>21</sup> | *Enterococcus faecalis* | *E. faecalis* PAI | $4.00 \times 10^{-10}$ | Also includes a HGT step |
| Poyart et al., 1995 <sup>20</sup> | *Enterococcus faecalis* | Tn916 | $1.48 \times 10^{-4}$ | Also includes a HGT step |
| Poyart et al., 1995 <sup>20</sup> | *Enterococcus faecalis* | Tn916 | $5.67 \times 10^{-9}$ | Also includes a HGT step |
| Poyart et al., 1995 <sup>20</sup> | *Enterococcus faecalis* | Tn916 | $2.40 \times 10^{-9}$ | Also includes a HGT step |
| Scott et al., 1988 <sup>18</sup> | *Bacillus subtilis* | Tn916 | $1.26 \times 10^{-9}$ | Also includes a HGT step |
| Franke and Clewell, 1981 <sup>17</sup> | *Streptococcus faecalis* | Tn916 | $9.95 \times 10^{-9}$ | Also includes a HGT step |

**Table S2.** Literature survey: Conjugation.

| Study | Species | Test system | Events/Day (average) | Remark |
| --- | --- | --- | --- | --- |
| Hausner and Wuertz, 1999 <sup>52</sup> | *Escherichia coli* | pRK415 | $5.76 \times 10^{-2}$ | |
| Wan et al., 2011 <sup>53</sup> | *Escherichia coli* | F-plasmid | $2.16 \times 10^2$ | |
| Simonsen et al., 1990 <sup>54</sup> | *Escherichia coli* | IncFII-plasmid R1 | $2.40 \times 10^{-10}$ | |
| Andrup and Andersen, 1999 <sup>55</sup> | *Escherichia coli* | F-plasmid | $2.16 \times 10^2$ | |
| Andrup and Andersen, 1999 <sup>55</sup> | *Enterococcus sp.* | pCF10 | $4.18 \times 10^2$ | |
| Zhong et al., 2012 <sup>56</sup> | *Escherichia coli* | pB10 | $3.12 \times 10^{-2}$ | |
| Jutkina et al., 2016 <sup>41</sup> | *Escherichia coli* | MGEs derived from wastewater | $3.84 \times 10^{-5}$ | Stress: Sub-MIC tetracycline concentration |
| Normander et al., 1998 <sup>57</sup> | *Pseudomonas putida* | TOL-plasmid | $4.46 \times 10^{-3}$ | |
| Suhartono and Sarvin, 2016 <sup>58</sup> | *Escherichia coli* | Plasmids derived from river water | $1.60 \times 10^{-3}$ | Stress: 20 mg/L nalidixic acid and 80 mg/L sulfamethoxazole or 20 mg/L streptomycin and 16 mg/L tetracycline |

**Table S3.** Dependency of the modelled processes on the MH and H parameters.

| Origin | MH:10<sup>-15</sup> to 10<sup>-13</sup> | MH:10<sup>-13</sup> to 10<sup>-10</sup> | MH:10<sup>-10</sup> to 10<sup>-7</sup> | MH:10<sup>-7</sup> to 10<sup>-4</sup> | MH:10<sup>-4</sup> to 10<sup>-2</sup> |
| --- | --- | --- | --- | --- | --- |
| pathogens in the human microbiome | 26.7% | 21.7% | 0.588% | 0.00194% | <0.00001% |
| non-pathogenic human-associated bacteria | 0.00659% | 1.56% | 36.3% | 83.4% | 96.0% |
| chromosomes of environmental bacteria | 0.00030% | 0.0633% | 1.38% | 3.65% | 2.92% |
| MGEs in human-associated bacteria | 70.1% | 73.7% | 59.4% | 12.4% | 1.02% |
| MGEs in the environment | 3.22% | 2.98% | 2.25% | 0.545% | 0.0311% |
| pathogenic bacteria in the environment | <0.00001% | <0.00001% | <0.00001% | <0.00001% | <0.00001% |
| Origin | H:10<sup>-11</sup> to 10<sup>-9</sup> | H:10<sup>-9</sup> to 10<sup>-7</sup> | H:10<sup>-7</sup> to 10<sup>-5</sup> | H:10<sup>-5</sup> to 10<sup>-3</sup> | H:10<sup>-3</sup> to 10<sup>-1</sup> |
| pathogens in the human microbiome | 99.0% | 73.3% | 3.63% | 0.0394% | 0.00033% |
| non-pathogenic human-associated bacteria | 0.717% | 5.90% | 3.07% | 0.303% | 0.226% |
| chromosomes of environmental bacteria | 0.0326% | 0.246% | 0.115% | 0.0130% | 0.00087% |
| MGEs in human-associated bacteria | 0.231% | 19.7% | 89.8% | 95.6% | 96.3% |
| MGEs in the environment | 0.0105% | 0.822% | 3.36% | 4.09% | 3.72% |
| pathogenic bacteria in the environment | 0.00001% | 0.00001% | <0.00001% | <0.00001% | <0.00001% |

## Additional supplementary material

Code is available from here: https://microbiology.se/publ/resistance_emergence_model/code.zip

Full results of the pre-existing (main) model: https://microbiology.se/publ/resistance_emergence_model/preexisting_model.zip

Full results of the emergence model: https://microbiology.se/publ/resistance_emergence_model/emergence_model.zip

